# Reconstitution and use of highly active human CDK1:Cyclin-B:CKS1 complexes

**DOI:** 10.1101/2021.09.24.461690

**Authors:** Pim J. Huis in ’t Veld, Sabine Wohlgemuth, Carolin Koerner, Franziska Mueller, Petra Janning, Andrea Musacchio

**Affiliations:** Department of Mechanistic Cell Biology, Max Planck Institute of Molecular Physiology, Dortmund, Germany; Department of Chemical Biology, Max Planck Institute of Molecular Physiology, Dortmund, Germany; Centre for Medical Biotechnology, Faculty of Biology, University Duisburg-Essen, Universitätsstrasse 45141 Essen, Germany

## Abstract

As dividing cells transition into mitosis, hundreds of proteins are phosphorylated by a complex of cyclin-dependent kinase 1 (CDK1) and Cyclin-B, often at multiple sites. CDK1:Cyclin-B phosphorylation patterns alter conformations, interaction partners, and enzymatic activities and need to be recapitulated *in vitro* for the structural and functional characterization of the mitotic protein machinery. This requires a pure and active recombinant kinase complex. The kinase activity of CDK1 critically depends on the phosphorylation of a Threonine residue in its activation loop by a CDK1 activating kinase (CAK). We developed protocols to activate CDK1:Cyclin-B either *in vitro* with purified CDK1 activating kinases (CAK) or in insect cells through CDK-CAK co-expression. To boost kinase processivity, we reconstituted a tripartite complex consisting of CDK1, Cyclin-B, and CKS1. In this work, we provide and compare detailed protocols to obtain and use highly active CDK1:Cyclin-B (CC) and CDK1:Cyclin-B:CKS1 (CCC).

## Introduction

A human cell that sets out to divide needs to condense DNA into chromosomes, break down the nuclear envelope, and form a mitotic spindle. Key for the initiation of these transformations is the rise in the activity of a complex between <u>C</u>yclin-<u>d</u>ependent <u>k</u>inase 1 (CDK1) and Cyclin-B, the main mitotic cyclin in vertebrates (Morgan, 2007; Hochegger et al., 2008). This results in the post-translational modification of hundreds of proteins, including other mitotic kinases, on specific serine and threonine side-chains and often at multiple sites (Daub et al., 2008; Olsen et al., 2010; Hegemann et al., 2011). Tight spatiotemporal regulation of these substrates and of their role in mitosis is then enabled by the intricate balance between mitotic kinase activities and counteracting phosphatases (Gelens et al., 2018).

Phosphorylation regulates the conformation, the activity, and the binding partners of the mitotic protein machinery. CDK1 is primed to initiate mitosis after Cyclin-B replaces Cyclin-A as the major CDK1-associated Cyclin and after the side-chains of Thr14 and Tyr15 in CDK1 are dephosphorylated. This is promoted by a shift in balance between the kinases Wee1 and Myt1 and the Cdc25 phosphatase, and ultimately discharges the glycine-rich loop (GEGTYG) from antagonizing substrate-binding and ATP hydrolysis (Morgan, 2007; Wood and Endicott, 2018). The activity of CDK kinases also critically depends on the configuration of its activation loop. The phosphorylation of this loop, at position Thr161 in CDK1, is required to dock the activation loop at the base of the active site, enabling efficient substrate binding and phosphate transfer (Jeffrey et al., 1995; Russo et al., 1996; Brown et al., 2015) (**Figure 1A, 1B**). Active CDK1 is thus phosphorylated at Thr161 and dephosphorylated at positions Thr14 and Tyr15.

**Figure 1:**
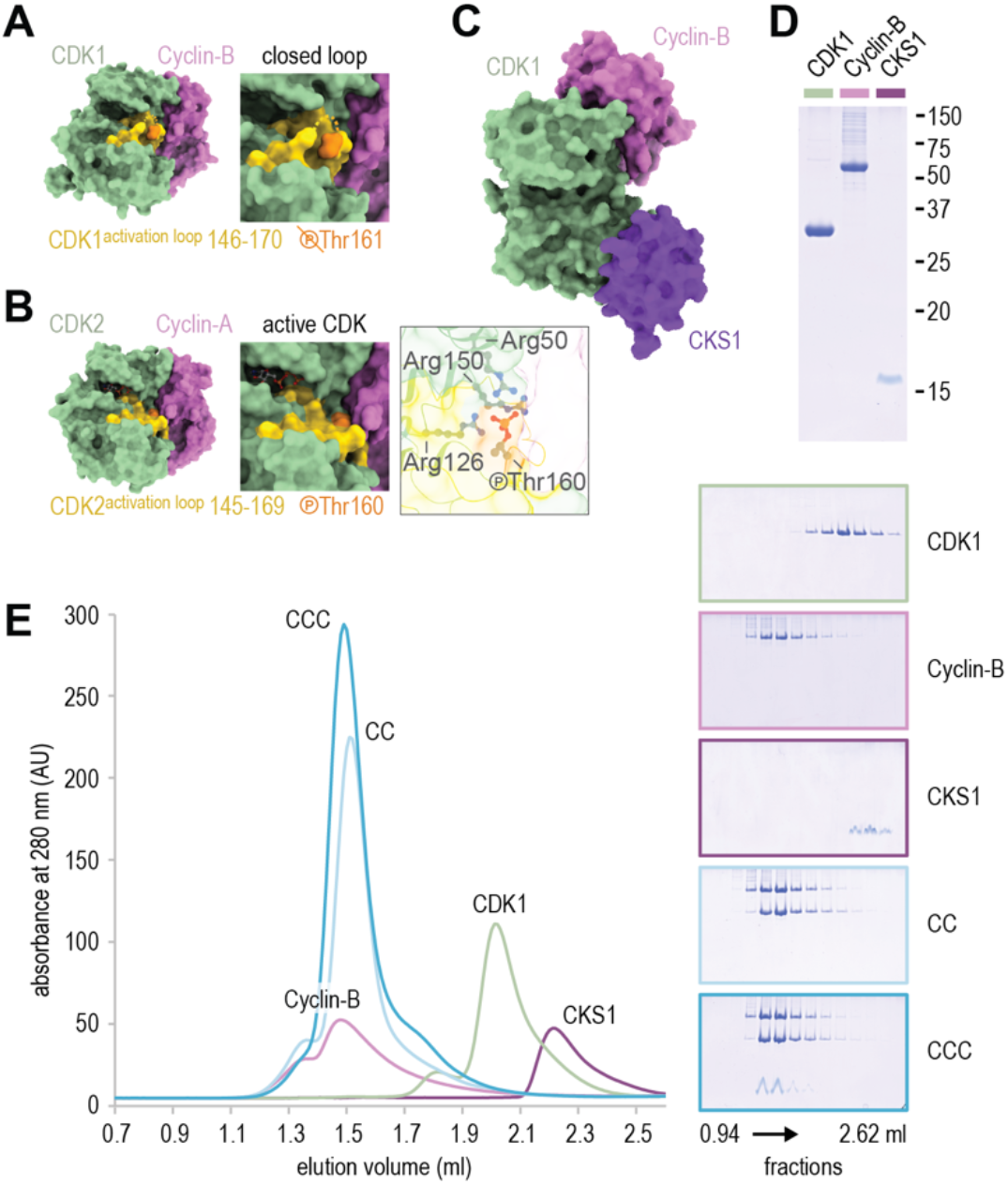
Reconstitution of stoichiometric CDK1:Cyclin-B and CDK1:Cyclin-B:CKS1 complexes. **a**) Surface view of a complex between CDK1 (green) and Cyclin-B (purple) (PDB 4YC3)(Brown et al., 2015). The CDK1^162-173^ activation loop is shown in gold and Threonine 161 in orange. Within the closed activation loop, residues CDK1^162-164^ (HEV) are not visible and replaced with a dashed line. **b**) Surface view of CDK2:Cyclin-A with bound ATP (PDB 1JST)(Russo et al., 1996). The side chains of Arginines 50, 126, and 150 coordinate the activation loop with a phosphorylated Threonine 160. **c**) Surface view of a tripartite CDK1:Cyclin-B:CKS1 complex (PDB 4YC3). Compared to panel a), the structure is rotated 45° along the x-axis. **d, e**) Analysis of purified CDK1, Cyclin-B, CKS1, CDK1:Cyclin-B (CC), and CDK1:Cyclin-B:CKS1 (CCC) by SDS-PAGE followed by Coomassie staining and size exclusion chromatography using a Superdex 200 increase 5/150 column.

Phosphorylation of the activation loop of CDK1 occurs in *trans* by a <u>C</u>DK-<u>a</u>ctivating <u>k</u>inase (CAK). In metazoans, CDK1 is activated by the CDK7:Cyclin-H kinase(Fisher and Morgan, 1994; Larochelle et al., 1998), which is also implicated, as a module of TFIIH, in the phosphorylation of the C-terminal domain of RNA polymerase II, as recently depicted (Greber et al., 2020; Abdella et al., 2021; Chen et al., 2021). Whereas metazoan CAK consists of a CDK7:Cyclin-H complex, activation of the budding yeast CDK1 (Cdc28) is mediated by the distantly related and monomeric CAK1 (hereafter scCAK1)(Espinoza et al., 1996; Kaldis et al., 1996; Thuret et al., 1996).

Besides phosphorylation, the CDK1:Cyclin-B kinase activity and substrate binding preferences are also regulated by various binding proteins. An important regulator is CKS1, a small adaptor protein that binds to the large helical lobe of the CDK1 kinase fold (**Figure 1C**). CKS1 contains a binding pocket for phosphorylated threonine residues and guides substrates with proximal phosphothreonine residues to the active site of CDK1, resulting in multisite phosphorylation (Kõivomägi et al., 2011, 2013).

With a persistent interest in the functional and structural characterization of proteins that act in cell division, and attempts to reconstitute *in vitro* fundamental aspects of the cell division process, it is essential to recapitulate CDK1:Cyclin-B phosphorylation patterns *in vitro* with a recombinant kinase complex that is pure, active, and processive. We have developed and compared ways to activate recombinant CDK1 with yeast or human CAK *in vitro* or in insect cells during protein expression. To boost the processivity of reconstituted CDK1:Cyclin-B complexes, we also set out to include CKS1 in our recombinant kinase complexes. Taken together, our analysis demonstrates how to reconstitute CDK1:Cyclin-B (CC) and CDK1:Cyclin-B:CKS1 (CCC) complexes, how to phosphorylate the activation loop of CDK1 efficiently, and how to reconstitute multi-site CDK1 phosphorylation patterns *in vitro*.

## Results

### Reconstitution of stoichiometric CDK1:Cyclin-B and CDK1:Cyclin-B:CKS1 complexes

We set out to reconstitute CDK1:Cyclin-B (CC) and CDK1:Cyclin-B:CKS1 (CCC) complexes from their purified individual constituents. We therefore expressed full-length CDK1, Cyclin-B1, and CKS1 in insect cells and purified these proteins to homogeneity using size-exclusion chromatography after affinity tags were proteolytically removed (**Figure 1D**). Stable and stoichiometric CC (83 kDa) and CCC (93 kDa) complexes formed when CDK1, Cyclin-B, and CKS1 were mixed and were further purified using size-exclusion chromatography (**Figure 1E**). These results highlights that the formation of CDK1:Cyclin-B or CDK1:Cyclin-B:CKS1 complexes does not require the phosphorylation of the CDK1 activation loop (see below).

### Phosphorylation of CDK1 in insect cells and *in vitro* by a CDK activating kinase

Since the phosphorylation of CDK1^Thr161^ by the <u>C</u>DK <u>a</u>ctivating <u>k</u>inase (CAK) promotes substrate binding and phosphorylation, we set out to co-express CDK1 and CAK in insect cells. Purified CDK1 that was co-expressed with either yeast or human CAK (scCAK1 or hsCDK7, respectively) was effectively phosphorylated, as judged by its retarded mobility on a phostag-containing acrylamide gel (Kinoshita et al., 2006) (**Figure 2**, lanes 1-3). We also purified scCAK1 (39 kDa) and used it to phosphorylate CDK1 *in vitro*. Analysis by phostag-PAGE showed effective phosphorylation of CDK1 in a reaction of 100 minutes at room temperature (**Figure 2**, lanes 4-8). CDK1 associated with Cyclin-B is also a CAK substrate, but the rate of phosphorylation was slower when CDK1 was in the Cyclin-bound state (**Figure 2**, lanes 4-13). The similar migration of Cyclin-B throughout the course of the experiment confirms that Cyclin-B is neither a CDK nor a CAK substrate.

**Figure 2:**
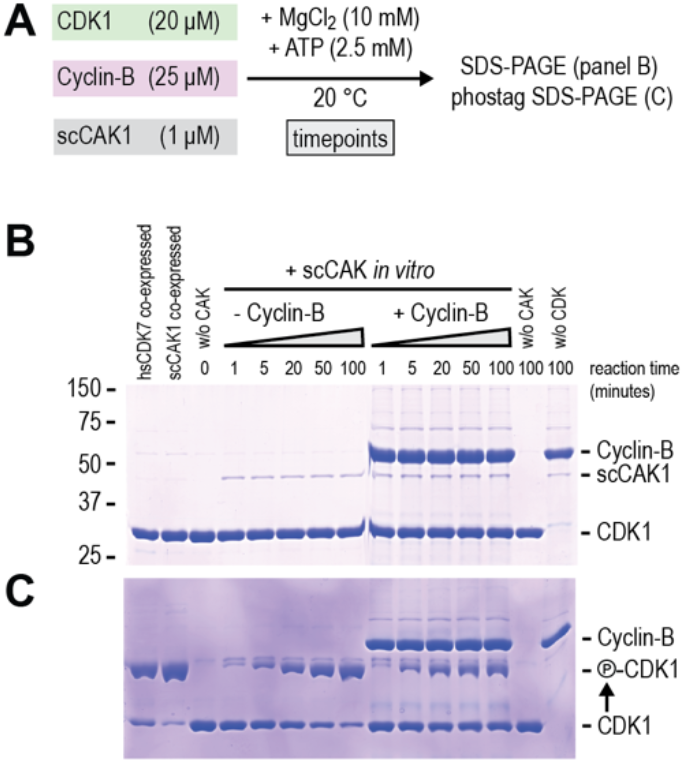
Phosphorylation of CDK1 in insect cells and *in vitro* by a CDK activating kinase. **a**) Reaction scheme for the phosphorylation of CDK1 with scCAK1. **b, c**) SDS-PAGE analysis of CDK1, Cyclin-B, and scCAK1. The presence of phostag-acrylamide (panel c) slows the migration of phosphorylated CDK1.

### Phosphorylation of CDK1 Threonine 161 activates recombinant CDK1:Cyclin-B

We next used semi-quantitative mass spectrometry to assess if the CAK-induced shift in CDK’s phostag-PAGE migration reflected phosphorylation of Threonine 161. Analysis of CDK1 that was exposed to human or yeast CAK in insect cells or *in vitro* confirmed the efficient phosphorylation of Threonine 161 and demonstrated that very few additional CDK1 residues, if any, were phosphorylated, and grossly substoichiometrically (**Supplementary Table 1**). Exposure to lambda-phosphatase reversed Thr161 phosphorylation (**Supplementary Figure 1**). The overall degree of Thr161 phosphorylation detected by mass spectrometry matches the intensities of the modified and unmodified CDK on phostag-PAGE (**Figure 3A-D, Supplementary Figure 1**). Whereas CDK1 was efficiently phosphorylated by scCAK1 both *in vitro* (92%) and upon co-expression in insect cells (91%), the phosphorylation by hsCDK7 was markedly lower *in vitro* than in insect cells (22% vs 78%) (**Figure 3A-D**, compare reactions 2, 3, 5, 7, and **Supplementary Figure 1**).

**Figure 3:**
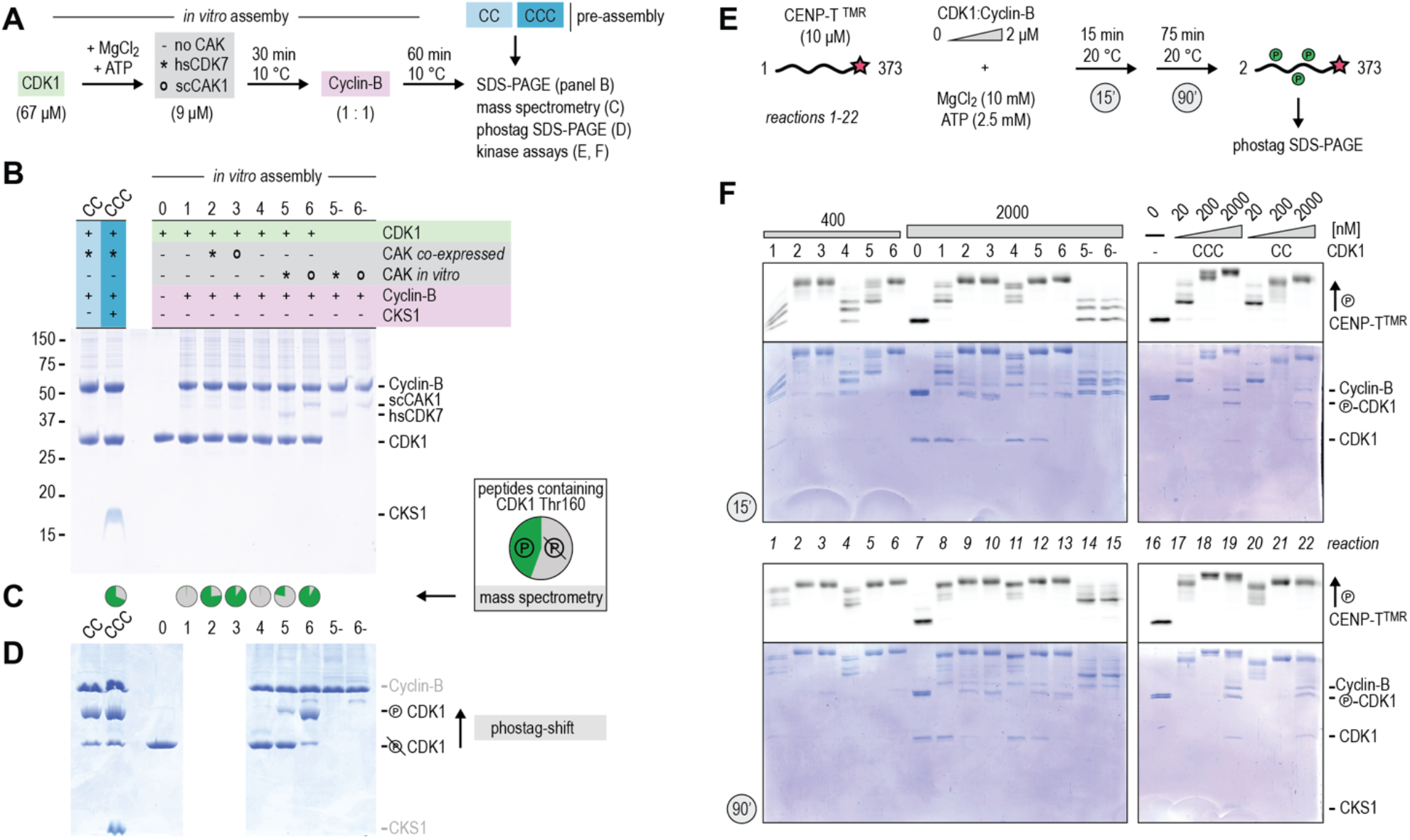
Phosphorylation of CDK1 Threonine 161 activates recombinant CDK1:Cyclin-B. **a, b**) Preparation and SDS-PAGE analysis of CDK1:Cyclin-B complexes that were either assembled from purified components or pre-assembled into dimeric or trimeric complexes. Reactions 1 and 4 are technical replicates. **c, d**) Selected CDK1:Cyclin-B complexes were analyzed for phosphorylation by mass spectrometry (technical triplicates) and phostag SDS-PAGE. Chart diagrams show the summed average intensities of all phosphorylated (green) and non-phosphorylated (grey) peptides containing Thr161. See **Supplementary figure 1** for more information. The effect of scCAK1 or hsCDK7 co-expression on the phostag-migration of CDK1 is shown in **Figure 2c**, lanes 1-3. **e, f**) Fluorescently labeled CENP-T was exposed to different kinase complexes and analysed for multi-site phosphorylation using phostag SDS-PAGE. The same gels were analysed for in-gel fluorescence (CENP-T) and Coomassie staining (all proteins). Reactions 1 and 4 as well as 8 and 11 are technical replicates.

To test and compare the activity of CDK1:Cyclin-B complexes, we used a 373-residue fragment encompassing the N-terminal region of the kinetochore protein CENP-T. This is an ideal multi-site phosphorylation model substrate as it is predominantly disordered and is phosphorylated by CDK1:Cyclin-B on multiple sites *in vivo* and *in vitro* (Nishino et al., 2013; Kim and Yu, 2015; Rago et al., 2015; Huis in ’t Veld et al., 2016). CENP-T^1-373^, expressed in bacterial and therefore unphosphorylated, was initially purified, labelled at its carboxy-end with tetramethylrhodamine (TMR) using Sortase-mediated conjugation (Guimaraes et al., 2013; Hirakawa et al., 2015), and finally purified to homogeneity using size-exclusion chromatography.

We used CENP-T at a concentration of 10 μM and varied the concentration of the CDK1:Cyclin-B complexes between 20 nM and 2 μM. To monitor its phosphorylation, we used in-gel fluorescence after phostag-PAGE (**Figure 3E, F**). CENP-T was not phosphorylated after 90 minutes in the absence of kinase or when incubated with CDK1 without Cyclin-B, highlighting the purity of the CENP-T and CDK1 samples and the strict Cyclin-dependency of CDK1 activity (**Figure 3F**, reactions 16 and 7). Sparse CENP-T phosphorylation occurred in the absence of CDK1, probably caused by traces of kinase in the Cyclin-B sample (**Figure 3F**, reactions 14 and 15). Relative to this control sample, phosphorylation appeared enhanced when CENP-T was incubated with CDK1:Cyclin-B that had not been exposed to a CAK. At CDK1:Cyclin-B concentrations of 0.4 or 2 μM, CENP-T underwent a progressive increase in phosphorylation between 15 and 90 minutes (**Figure 3F**, reactions 1, 4, 8, and 11). By contrast, apparently homogeneous multi-site CENP-T phosphorylation patterns were already visible after 15 minutes of phosphorylation with 0.2 or 0.4 μM CDK1:Cyclin-B complexes generated with kinase co-expressed with scCAK1 or hsCDK7 in insect cells (**Figure 3F**, reactions 2, 3, and 21). After a 90-minute reaction, comparable CENP-T phosphorylation levels were observed with concentrations of 20 nM or 2 μM of CAK-treated or CAK-untreated CDK1:Cyclin-B complexes, respectively (**Figure 3F**, reactions 8, 11, and 20). CAK exposure thus increases the activity of recombinant CDK1:Cyclin-B complexes on CENP-T by approximately two orders of magnitude. Consistent with a modest but marked increase in the phosphorylation levels of CDK1 Thr161, the *in vitro* activation of CDK1:Cyclin-B with hsCDK7 resulted in increased CENP-T phosphorylation (**Figure 3F**, reactions 4 and 5). However, CDK1 activation *in vitro* with scCAK1 was more effective (**Figure 3F**, reactions 5 and 6), indicating that the phosphorylation of Thr161 correlates with kinase activity.

### CKS1 boosts the processivity of CDK1:Cyclin-B

A side-by-side comparison of pre-assembled CDK1:Cyclin-B (CC) and CDK1:Cyclin-B:CKS1 (CCC) complexes suggested the existence of a slower migrating form of CENP-T in the CCC reactions (**Figure 3F**, reactions 18, 19, 21, and 22). To investigate putative effects of CKS1 on CDK1:Cyclin-B processivity, we followed the phosphorylation of CENP-T with 100 nM of CC or CCC over time (**Figure 4A, B**). These complexes contain CDK1 that was activated by scCAK1 co-expression and were reconstituted from individual constituents with (pre-assembled) or without (*in vitro* assembly) size-exclusion chromatography (**Figure 4A, B**).

**Figure 4:**
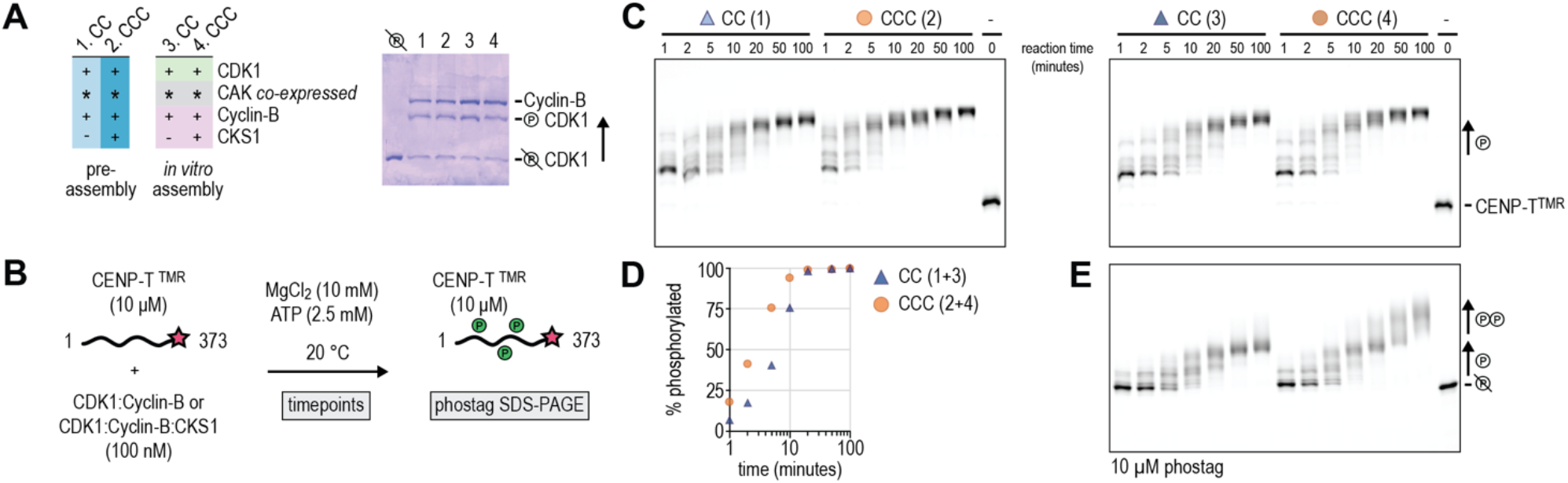
CKS1 boosts the processivity of CDK1:Cyclin-B. **a**) Preparation and phostag SDS-PAGE analysis of CDK1:Cyclin-B and CDK1:Cyclin-B:CKS1 complexes that were either assembled from purified components or pre-assembled into dimeric or trimeric complexes. **b, c**) CENP-T was exposed to different kinase complexes and analysed for multi-site phosphorylation using phostag SDS-PAGE and in-gel fluorescence (CENP-T^TMR^). **d**) Quantification of phosphorylated / non-phosphorylated signals from the gels shown in panel c). **e**) SDS-PAGE analysis of samples CC (3) and CCC (4) as in panel c) in the presence of 10 μM phostag acrylamide (all other gels contain 50 μM phostag-acrylamide).

Following these reactions over time revealed that the presence of CKS1 sped up CENP-T phosphorylation approximately two-fold (**Figure 4C, D**). In addition to faster phosphorylation, CENP-T migration appeared to continuously decrease over time in the presence of CCC. This suggests that low-affinity CENP-T sites become CDK1 substrates when proximal phosphorylated threonine residues dock to CKS1 (Kõivomägi et al., 2011, 2013). The difference between high (∼5-10 sites) and hyper (10+ sites) phosphorylation on CENP-T was most apparent when the phostag concentration in the polyacrylamide gel was lowered five-fold (**Figure 4E**). Taken together, we demonstrate that the presence of CKS1 enables CDK1:Cyclin-B to efficiently establish multi-site phosphorylation patterns.

## Discussion

Over the last decades, genetic, biochemical, and structural studies provided detailed insights into the regulation of the cell cycle by CDK:Cyclin combinations and the activation of Cyclin-dependent Kinases by CAKs (Morgan, 2007; Wood and Endicott, 2018). Since hundreds of CDK:Cyclin phosphorylation events are required to rewire the protein machinery in dividing mitotic or meiotic cells, *in vitro* reconstitution of phosphorylation patterns for the functional and structural characterization of simplified, smaller subsets of components is mandatory. To facilitate this type of experiments, the availability of pure and active recombinant CDK1:Cyclin-B complexes is essential, and here we have contributed to the development of a strong pipeline towards this goal. A previous study demonstrated how to obtain a pure and active CDK1:Cyclin-B complex using purified scCAK1 (expressed in bacteria) and CDK1 (expressed in insect cells), *in vitro* phosphorylation of CDK1 with scCAK1, and subsequent assembly of the active CDK1:Cyclin-B complex (Brown et al., 2015). In this study, we built on this previous work and took a systematic approach to optimize protocols for the generation of highly active and processive recombinant CDK1:Cyclin-B:CKS1 complexes. These robust and easy-to-implement protocols and the corresponding expression plasmids, publicly available through Addgene (https://www.addgene.org/), will be a valuable resource for researchers with a range of backgrounds and an interest in investigating the protein machinery orchestrating cell division. We demonstrate how in-gel fluorescence and general protein staining after phostag-PAGE is a cost-effective and swift combination to analyze single and multi-site phosphorylation. Whereas mass spectrometry and site-directed mutagenesis of putative phosphorylation sites are needed to specify modified residues, phostag-PAGE provides a clear overview of phosphorylation coverage -for example to follow the amounts of phosphorylated and non-phosphorylated CDK1 Thr161- and can readily be combined with immunofluorescence. The latter is especially useful to study more complex reaction mixtures.

Co-expression of CDK1, Cyclin-B, and CKS1 in insect cells was used in a very recent study that revealed how recombinant CCC complexes form a stoichiometric complex with Separase (Yu et al., 2021). In that study, the CCC assembly complexed to Separase had been obtained by insect cell co-expression, but had not been co-expressed with CAK or treated with CAK activity during purification. Surprisingly, however, density for the phosphorylated Thr161 residues of CDK1 was clearly visible, demonstrating that at least a significant subset of CDK1 might have been targeted by an endogenous CAK activity in insect cells. This may seem to imply that monomeric CDK1, which we find to be very little (<1%) phosphorylated on Thr161 when expressed as an isolated subunit without exogenous co-expression of CAK, is not a good substrate of the endogenous CAK activity of insect cells. This is consistent with previous studies describing that CDK7:Cyclin-H can phosphorylate CDK2 but not CDK1 in their Cyclin-free forms (Fisher and Morgan, 1994).

Our finding that hsCDK7 is a potent CDK1 activator when co-expressed in insect cells but does not phosphorylate CDK1 efficiently *in vitro* also suggests that host-cell (ovary cells from the cabbage looper moth *Trichopulsia ni*) factors contribute to CAK activation that leads to the phosphorylation of CDK1’s Thr161 (**Figure 3**). Such factors could include a hsCDK7 activating kinase (CAKAK) or a Cyclin that associates with hsCDK7 or CDK1. In fission yeast, it was for example demonstrated that the kinase Csk1 acts as a CAK and as a CAKAK (Hermand et al., 1998; Saiz and Fisher, 2002). In our work, we maximized robustness and yields by reconstituting CC or CCC complexes after individual expression of the constituents, and CDK1 was specifically activated through exposure to scCAK1 during co-expression or *in vitro*.

How multi-site phosphorylation rewires proteins that govern cell division was highlighted in a study showing that reconstituted APC/C could only bind its activator CDC20 after a region in the APC1 subunit was hyperphosphorylated and released from a CDC20 binding site on APC8 (Zhang et al., 2016). In that work, APC/C was phosphorylated *in vitro* with PLK1 kinase and a CDK2:Cyclin-A:CKS2 complex in a manner that reflects APC/C phosphorylation in mitosis (Kraft et al., 2003; Hegemann et al., 2011; Zhang et al., 2016). Although the constituents of the CDK2:Cyclin-A:CKS2 complex were individually expressed in bacteria and had not been exposed to a CAK, and were therefore most probably not phosphorylated, the resulting kinase did effectively reconstitute multi-site phosphorylation patterns on APC/C. CDK2:Cyclin-A:CKS2 has been previously shown to be strongly dependent on the phosphorylation of the activation loop in CDK2 (Hagopian et al., 2001). Thus, we speculate that the docking of CDK2:Cyclin-A:CKS2 on APC/C results in CAK-independent substrate phosphorylation. Supporting the latter explanation, we see that CDK1:Cyclin-B that had not been exposed to CAK efficiently phosphorylated CENP-T at concentrations of 0.4 and 2 μM in 90 minutes (**Figure 3**). Accordingly, we have used CDK1:Cyclin-B complexes that were not CAK-exposed at relatively high concentrations of 0.25 - 1 μM in recent studies to successfully phosphorylate substrates using reactions for 16 hours at 4ºC (Huis in ‘t Veld et al., 2019; Piano et al., 2021; Singh et al., 2021).

In summary, we describe the discrete effects of activation loop phosphorylation and CKS binding on the activity and processivity of a recombinant CDK1:Cyclin B complex, together with protocols for expression and access to the relevant expression vectors. These tools will support work of *in vitro* reconstitution of crucial aspects of cell division.

## Materials and Methods

### Baculovirus generation and protein expression in insect cells

CDK1, Cyclin-B1, CKS1B, and hsCDK7 cDNA constructs were codon-optimized for insect cell expression and obtained from GeneArt (Life Technologies). The scCAK1 (CIV1) construct was kindly provided by David Barford. Expression cassettes of CDK1, Cyclin-B, CKS1B, scCAK1, and hsCDK7 were inserted into pLIB vectors (Weissmann et al., 2016) with N-terminal GST-3C (CDK1) or Polyhistide-TEV (others) tags and baculoviruses were generated following standard protocols (Trowitzsch et al., 2010). For expression, *Sf9* cells were infected for 3-4 days and added (1:20 dilution) to logarithmically growing *Hi5*-derived *Tnao38* insect cells (Hashimoto et al., 2012). For co-expressions, *Sf9* cultures expressing CDK1 (1:20 dilution) and scCAK1 or hsCDK7 (1:40 dilution) were simultaneously added to the expression culture. All insect cells were kept at 27°C. Cells were harvested after 3 days by centrifugation at 1000*g* at room temperature, washed once with ice-cold PBS, pelleted by centrifugation at 1000*g* at 4 °C, flash-frozen in liquid nitrogen, and stored at -80 °C.

### CDK1 purification

All protein purification steps were performed on ice or at 4 °C. Pellets from 2 liters of insect cells expressing GST-3C-CDK1 (with or without CAK co-expression) were resuspended in 160 ml buffer A (50mM HEPES pH7.4, 250mNaCl, 2mM TCEP, 5% v/v glycerol) supplemented with HP plus protease inhibitor mix (Serva) and DNase I (Roche). Cell lysates were prepared by sonication and cleared by centrifugation at 80000g at 4°C for 30-45 minutes. The soluble lysate was passed through a 0,8 μm filter and loaded onto a column with 20 ml Gluathione Sepharose 4 Fast Flow resin (Cytiva). After washing with 25 column volumes of buffer A, CDK1 was cleaved with in-house generated 3C protease for 16 hours. The eluate was concentrated to 2 ml through centrifugation with a 30k Amicon filter (Millipore) and separated on a Superdex 200 16/600 column equilibrated in buffer A. To remove GST, uncleaved GST-CDK1, and GST-3C-PreScission, a 5 ml GSH column (GE Healthcare) was mounted after the size-exclusion column. Selected fractions were concentrated to concentrations well above 100 μM. Purification from 2l of insect cells yielded approximately 4 mg of CDK1.

### Cyclin-B purification

His-TEV-Cyclin-B containing lysates were prepared as described above with 15 mM imidazole in lysis and wash buffers. After loading onto a 5ml or 10 ml Talon (Clontech) or Ni-NTA (GE Healthcare) column, and washing with approximately 50 column volumes, Cyclin-B was eluted in buffer A with 250 mM imidazole and concentrated to 2 ml through centrifugation with a 30k Amicon filter (Millipore). To remove the polyhistidine tag, Cyclin-B was exposed to TEV protease for 16 hours. Cyclin-B was further purified on a Superdex 200 16/600 column equilibrated in buffer A. To remove His-TEV protease and uncleaved His-Cyclin-B, a 5 ml Talon column (GE Healthcare) was mounted after the size-exclusion column. Selected fractions were concentrated to concentrations well above 100 μM. Purification from 1l of insect cells yielded approximately 15 mg of Cyclin-B.

### CKS1 purification

His-TEV-CKS1 was purified as described for Cyclin-B but using 5k Amicon filters and a Superdex 75 16/600 size-exclusion column. Approximately 5 mg of CKS1 was obtained from 0.5l of insect cells.

### CDK1:Cyclin-B (CC) and CDK1:Cyclin-B:CKS1 (CCC) formation

Purified CDK1, Cyclin-B, and CKS1 were mixed in a 1 : 1 : 0 (CC) or 1 : 1 : 2 (CCC) ratio for 1-2 hours on ice with concentrations above 100 μM. After size-exclusion chromatography on a Superdex 200 16/600 column in buffer A, fractions containing CC or CCC were concentrated through centrifugation with a 30k Amicon filter (Millipore), flash-frozen in liquid nitrogen, and stored at -80°C. Analytical size-exclusion chromatography (Figure 1) was performed using a Superdex 200 5/150 column.

### scCAK1 and hsCDK7 purification

The CDK1 activating kinases scCAK1 and hsCDK7 were purified as described for Cyclin-B above and yielded approximately 0.5 mg and 5 mg, respectively.

### CENP-T purification

CENP-T^2-373^ was expressed, purified, and fluorescently labelled as described (Huis in ‘t Veld et al., 2016). In brief, the expression of GST-3C-CENP-T^2-373^ with a C-terminal -LPETGG extension was induced in *E. coli* BL21(DE3)-codon-plus-RIL cells through the addition of 0.35 mM IPTG for ∼14 hours at 20°C. Subsequent steps were performed at 4°C or on ice. Cleared lysates were prepared by sonication and centrifugation and bound to a Glutathione-Agarose resin (Serva). GST-3C-CENP-T was cleaved off the beads with in-house generated 3C protease for 16 hours. After further purification using a Heparin HP column (GE Healthcare), CENP-T was fluorescently labeled with a GGGGK-TMR (5-Carboxytetramethylrhodamine) peptide (GenScript) using the Calcium-independent Sortase 7M (Hirakawa et al., 2015). After size-exclusion chromatography using a Superdex 200 16/600 column (GE Healthcare) in a buffer containing 20 mM Tris-HCl, pH 8.0, 150 mM NaCl, and 1 mM TCEP, CENP-T^2-373 TMR^ was concentrated to 118 μM and stored at -80°C.

### *In vitro* phosphorylation

Phosphorylation reactions were performed in the presence of MgCl_2_ (10 mM) and ATP (2.5 mM) and monitored on standard denaturing 10% polyacrylamide gels (Laemmli, 1970) in the presence of 50 μM phos-tag-acrylamide (Fujifilm Wako Chemicals)(Kinoshita et al., 2006). The *in vitro* phosphorylation of CDK1 by scCAK1 and hsCDK7 is described in detail in **Figure 2** and **Figure 3**. The *in vitro* phosphorylation of CENP-T^2-373 TMR^ by CDK1:Cyclin-B and CDK1:Cyclin-B:CKS1 complexes is described in detail in **Figure 3** and **Figure 4**.

### Mass spectrometry of CDK1

To analyze the phosphorylation of CDK1 by mass spectrometry, samples were reduced, alkylated and digested with LysC and Trypsin and prepared as previously described (Rappsilber et al., 2007). Obtained peptides were separated on an U3000 nanoHPLC system (Thermo Fisher Scientific). Samples were injected onto a desalting cartridge, desalted for 5 min using water in 0.1% formic acid, backflushed onto a Pepmap C18 nanoHPLC column (Thermo Fisher Scientific) and separated using a gradient from 5–30% acetonitrile with 0.1% formic acid and a flow rate of 300 nl/min. Samples were directly sprayed via a nano-electrospray source in an Orbitrap type mass spectrometer (Thermo Fisher Scientific). The mass spectrometer was operated in a data-dependent mode acquiring one survey scan and subsequently up to fifteen MS/MS scans. To identify phospho-sites, the resulting raw files were processed with MaxQuant (version 1.6.17.0) searching for CDK1, Cyclin-B, and CKS1 sequences with acetylation (N-term), oxidation (M) and phosphorylation (STY) as variable modifications and carbamidomethylation (C) as fixed modification. A false discovery rate cut off of 1% was applied at the peptide and protein levels and as well on the site decoy fraction (Cox and Mann, 2008).

### Structures

Surface views of CDK1:Cyclin-B:CKS1 (PDB 4YC3) (Brown et al., 2015)and of CDK2:Cyclin-A with bound ATP (PDB 1JST) (Russo et al., 1996) were prepared using Chimera X (version 0.9) (Goddard et al., 2018).

### Reagent Availability

The following plasmids are available through Addgene (addgene.org/Andrea_Musacchio/): pLIB-GST-3C-CDK1 (177011), pLIB-HIS-TEV-Cyclin-B (177012), pLIB-HIS-TEV-CKS1(177013), and pLIB-HIS-TEV-scCAK1(177014). All other reagents are available upon request.

## Author Contributions

CRediT contributorship: Conceptualization PH, AM; Formal Analysis PH; Funding acquisition AM; Investigation PH, FM, PJ; Methodology PH, SW, CK; Project Administration PH; Resources SW, CK; Supervision PH, AM; Visualization PH; Writing original draft PH; Writing – review & editing PH, AM.

## Acknowledgements

We thank David Barford for providing an scCAK1 expression plasmid, Priyanka Singh for early experiments with recombinant CDK1:Cyclin-B, and Kai Walstein for the preparation of TMR labeled CENP-T. AM gratefully acknowledges funding through the European Research Council Synergy Grant SyG 951430 BIOMECANET.

---

**Supplementary Figure 1:**
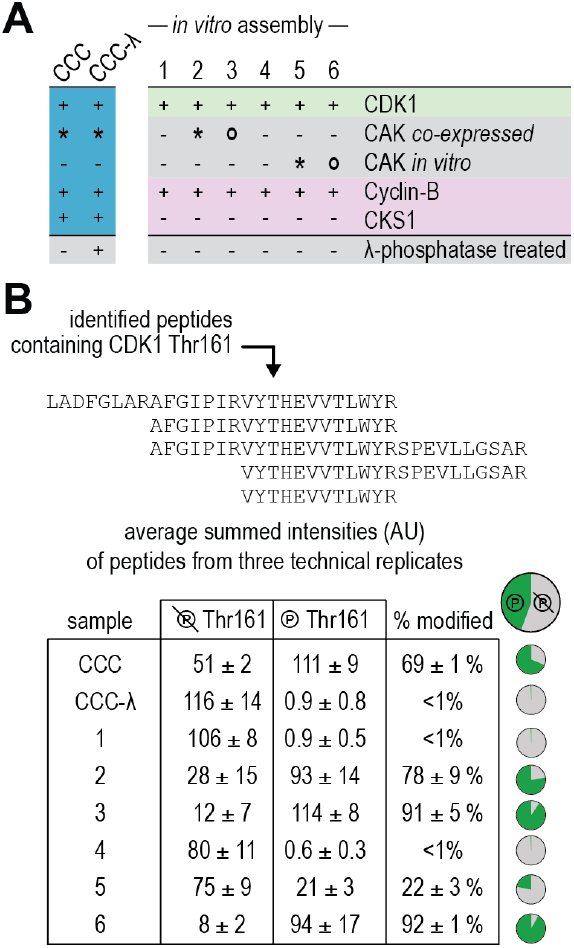
Analysis of phosphorylation of CDK1’s Threonine 160 by mass spectrometry. a) Overview of samples analyzed by mass spectrometry. With the exception of CCC-λ, all samples were also included in **Figure 3**. b) Five different peptides containing CDK1 Thr 161 were detected after trypsin digestion. From three technical replicates, the average summed intensities (AU) were added for peptides with phosphorylated Thr 161 and for peptides with non-phosphorylated Thr 161. Intensities of phosphopeptides as a fraction of the total peptide peptide intensities are shown in the last column.

**Supplementary Table 1:**
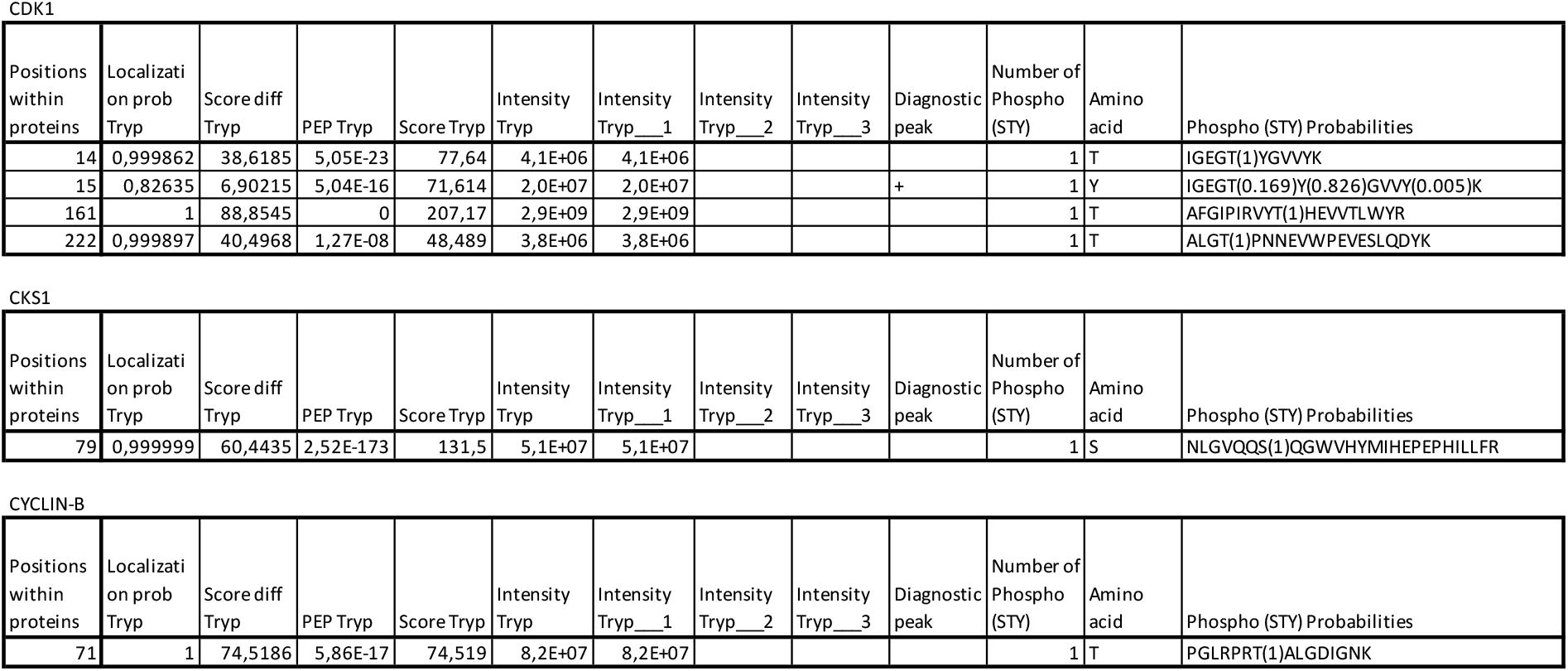
Phosphorylation in CDK1:Cyclin-B:CKS1. See **Figure 3** and **Supplementary Figure 1** for an in-depth analysis of the kinase activity of this sample (CCC) and the phosphorylation of Thr161 in CDK1.

